# Metabolic priming recruits tumour cells across a mitochondrial vulnerability threshold and sensitizes them to chemotherapy

**DOI:** 10.64898/2026.11.13.766410

**Authors:** Francisco J. Iborra, Raúl Guantes, Juan Díaz-Colunga, Antonio Martínez-Lorente, Ana Calatrava, José Luis Girela, Pedro M. Iborra, Juana Calabuig

**Author notes:** **Corresponding author.** Francisco J. Iborra, Instituto de Biomedicina de Valencia, IBV (CSIC), Jaume Roig 11, 46010 Valencia, Spain. ORCID 0000-0002-0692-3696.

## Abstract

Intratumour heterogeneity allows a fraction of cancer cells to escape otherwise effective treatment. Here we ask whether metabolic priming can recruit heterogeneous tumour cells into a transient mitochondrial state that is selectively vulnerable to photodynamic damage. Tetramethylrhodamine methyl ester (TMRM) has previously been shown to act as a mitochondria-targeted photosensitizer; we exploit its voltage-dependent accumulation and dynamic fluorescence during pulsed illumination to quantify vulnerability in single cells. The response is a reproducible fluorescence excursion, a rise to a maximum followed by a decay, accompanied by reactive oxygen species production. Comparing seventeen descriptors of this excursion against subsequent fate in 195 cells across four arms shows that quantities referenced to each cell’s own resting value outperform static intensity. The simple difference Fmax − F0 predicts death with held-out areas under the curve of approximately 0.75–0.79, whereas resting intensity alone carries little transferable information and time to peak does not discriminate fate in the datasets tested. Metabolic priming with the PKM2 activator TEPP-46 or with ethyl pyruvate increases the fraction of responding cells; under TEPP-46 almost the whole population enters the responding state. PKM2 abundance is associated with faster entry into that state, and PKM2 is heterogeneous in human skin tumour sections. Four hours of camptothecin shifts the response distribution, increasing the fraction of cells above the untreated median from 50% to 76%, identifying a candidate intervention window. In combination experiments, TMRM photodynamic treatment produced survival below the Bliss-independence expectation with camptothecin or TRAIL, in both treatment orders and across three cell lines; these combination measurements are descriptive and require independent replication. The study therefore identifies a perturbation-revealed mitochondrial vulnerability that can be pharmacologically recruited and followed dynamically at single-cell resolution. It is a proof of concept in cultured cells and human tumour sections and has not been tested in vivo.

**Highlights:**

- Pulsed illumination of TMRM reveals a reproducible mitochondrial response whose magnitude predicts subsequent cell fate.
- The excursion measured against each cell’s own resting signal, Fmax − F0, is more informative and more robust across experiments than resting intensity alone; time to peak does not discriminate fate in the datasets tested.
- PKM2 activation or pyruvate supplementation recruits otherwise non-responding cells into the photodynamically vulnerable state, so that under TEPP-46 almost the whole population responds.
- TMRM photodynamic treatment produces survival below the Bliss-independence expectation when combined with camptothecin or TRAIL in the experiments tested.
- PKM2 is heterogeneous in human skin tumour sections, supporting its evaluation as a candidate marker of metabolic response heterogeneity.

## Introduction

Resistance of tumour cells to antitumour therapy is one of the central problems of contemporary oncology [1,2]. It limits the efficacy of current treatments and it contributes to recurrence and metastasis, the leading causes of cancer mortality. Cells that survive therapy often acquire more aggressive phenotypes, which makes subsequent treatment less effective and leads to progression [3,4].

Therapeutic resistance can arise from pre-existing phenotypic variation, therapy-induced adaptation and their interaction. Heterogeneous tumours contain cells with different probabilities of surviving the same insult, while treatment itself can reorganise metabolism, signalling, stress responses and cell state. These processes are temporally interleaved rather than mutually exclusive, and the interval before residual states become stabilised may expose transient vulnerabilities [3,4,47]. Early intervention therefore matters not only because resistant cells may already exist, but because treatment continuously changes the distribution of states from which resistance can emerge.

Our laboratory has concentrated on the first hours of chemotherapy. Chemotherapy induces cellular stress, which triggers a cascade of adaptive responses. One event in that cascade is activation of Bax, which binds mitochondria and inhibits complex I of the electron transport chain. That inhibition raises mitochondrial membrane potential and increases production of reactive oxygen species (ROS). ROS act in two directions: at high levels they kill, at moderate levels they promote survival by activating pathways that support adaptation and proliferation [4]. Under elevated ROS, proteins are oxidatively damaged and degraded by the proteasome, intracellular amino acid concentrations rise, mTOR is activated and protein synthesis increases. We previously identified proteolysis and protein synthesis as critical to this response, and showed that targeting them alongside chemotherapy enhances tumour cell death [4].

A further feature associated with therapy adaptation is mitochondrial hyperpolarization. Lipophilic cationic fluorophores such as tetramethylrhodamine methyl ester (TMRM) accumulate in mitochondria in proportion to membrane potential. TMRM has already been shown to act as a mitochondria-targeted photosensitizer in glioblastoma cells [46]. Here we ask a different question: whether cell-to-cell metabolic heterogeneity determines entry into the TMRM-responsive state, whether that state can be pharmacologically recruited, and whether its dynamics can be used to identify a temporally exploitable vulnerability. This reframes TMRM-PDT from a photosensitizer alone into a perturbation that both reveals and acts on heterogeneous mitochondrial states.

Phenotypic heterogeneity produces cell-to-cell differences in mitochondrial content and dynamics [5,8–11], and those differences both cause differential resistance to cell death and drive metabolic reprogramming. This study is a proof of concept for attacking that process early, through a mitochondrially targeted approach.

## Results

### Pulsed illumination drives a reproducible excursion of TMRM emission

To characterize the dynamic response to TMRM photoactivation, we loaded HeLa cells with TMRM for 30 min and performed time-lapse fluorescence microscopy, acquiring an image every 10 s. In a fraction of cells the fluorescence of TMRM rose on irradiation until it reached a maximum and then began to fall (Figure 5a). We incubated cells for 30 min with TMRM from 6.25 nM to 2 µM and counted the cells that showed this excursion. The result is a concentration-dependent sigmoidal curve: below 250 nM no cells responded, whereas above 500 nM nearly all did (Figure 5b). The curve fits a Hill function with a coefficient of approximately 5, indicating a sharp threshold in the measured response. Previous work established that TMRM can act as a mitochondria-targeted photosensitizer [46]; the question here is why only some cells cross the dynamic response threshold and whether that state can be deliberately recruited.

A Hill coefficient of that size means that a small difference in effective mitochondrial dye concentration separates cells that show the excursion from those that do not. We treat the threshold as an empirical property of the response rather than as evidence for one specific photophysical mechanism. Self-quenching with subsequent dequenching, aggregation in the lipid environment and other concentration-dependent effects could contribute [12]. We do not have the measurements required to distinguish among these mechanisms. What matters for the rest of this work is that the excursion is reproducible, occurs only above a sharp response threshold, and contains information that can be related to subsequent cell fate.

The sub-mitochondrial environment of TMRM is relevant to its photodynamic behaviour, but voltage-driven accumulation does not by itself imply binding to a mitochondrial component. Förster resonance energy transfer (FRET) can report nanometre-scale proximity between fluorophores [13]. We therefore asked whether TMRM and MitoTracker Green (MG), which occupies the mitochondrial lipid environment, show reciprocal fluorescence dynamics consistent with close proximity during photoactivation.

MG can act as a fluorescence donor to TMRM when the two fluorophores are sufficiently close. In a classical acceptor-photobleaching experiment, loss of the acceptor releases donor quenching [14]. We co-stained cells with MG and TMRM and followed both signals during repeated illumination (Figure 5e,f). After aligning trajectories to the TMRM maximum, the onset of TMRM decay coincided with an abrupt increase in MG fluorescence. This reciprocal behaviour is consistent with loss of energy transfer as TMRM is depleted or photobleached and therefore with close proximity of the dyes in the mitochondrial membrane environment. Because the experiment does not provide a complete FRET calibration and alternative causes of reciprocal fluorescence cannot be excluded, we use it as supportive rather than definitive evidence of membrane association.

### The excursion is heterogeneous between cells and depends on membrane potential

Fluorescence enhancement appears only in some cells of a clonal population when TMRM is titrated at intermediate concentrations (Figure 5a,b). In responding cells the dynamics are a unimodal curve with three phases (Figure 1a, and see Figure 5e): an initial lag, a rapid rise and a maximum followed by a slower decay. The maximum marks the transition from accumulation/enhancement to signal loss and subsequent mitochondrial depolarization. Because loss of membrane potential often accompanies mitochondrial apoptotic commitment, this transition may occur near MOMP in dying cells, but the TMRM trajectory alone does not establish MOMP.

**Figure 1.**
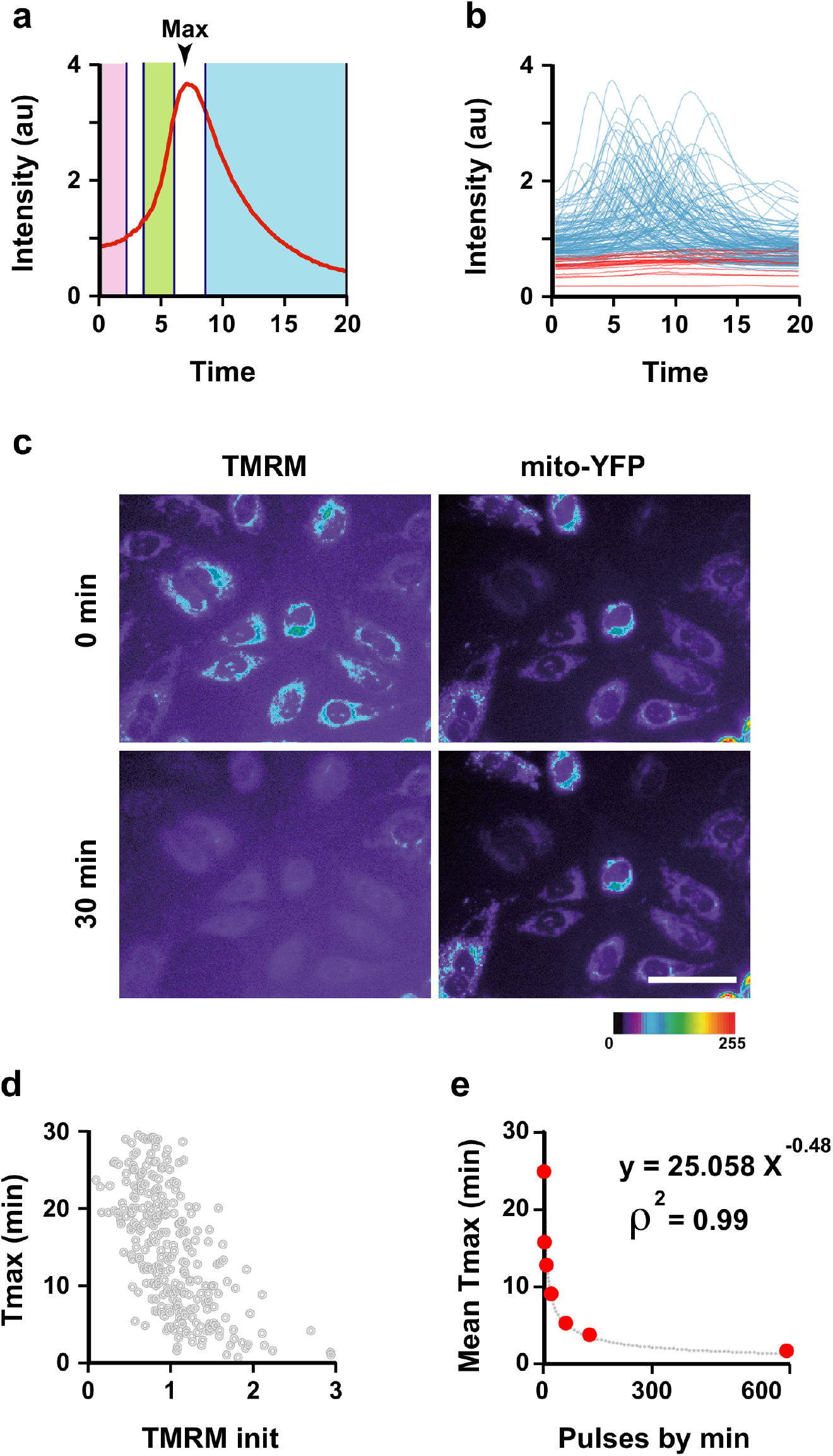
**Single-cell kinetic analysis**. (a) Integrated fluorescence of a single cell, with four features marked: a latency phase (pink), an exponential growth phase (green), the maximum, which marks the transition from fluorescence enhancement to signal loss, and a decay phase (blue). (b) Every cell follows the same phases with different kinetics, illustrating population heterogeneity. Blue, cells showing emission enhancement; red, cells not showing it. (c) TMRM signal loss does not coincide with gross loss of matrix integrity. TMRM and transfected mito-YFP, which localizes to the matrix, are shown at 0 and 30 min. After 30 min TMRM fluorescence has largely disappeared while mito-YFP remains localized. This rules out gross mitochondrial rupture but does not by itself establish MOMP. (d) Time to peak against initial integrated TMRM signal in individual cells: cells with lower initial intensity tend to reach the maximum later. (e) Tmax against light-pulse frequency at constant intensity per pulse. Tmax falls as pulse rate rises. Each rate was acquired in a separate session, so the fitted exponent is descriptive.

We asked whether the decline after the maximum reflected gross mitochondrial rupture. Cells were transfected with YFP-mito, which localizes to the matrix [16]. Disruption sufficient to release matrix contents would delocalize YFP-mito, whereas YFP-mito remained tightly localized throughout the TMRM decay (Figure 1c). The experiment therefore rules out gross structural rupture as the explanation for signal loss. It does not by itself distinguish mitochondrial depolarization from outer-membrane permeabilization, so throughout the manuscript we treat the excursion as a photodynamic mitochondrial response rather than as a direct measurement of MOMP.

Looking at individual cells, two features stand out. Peak times are highly variable (Figure 1b), as is common for mitochondrial and apoptotic responses in clonal populations [5,17]. Cells that show no enhancement also tend to have lower initial TMRM intensity (red traces, Figure 1b), consistent with lower initial membrane potential. Because initial TMRM fluorescence is tightly correlated with initial MG fluorescence, a proxy for mitochondrial content [11], and because mitochondrial content varies widely between cells [10,18], both mitochondrial mass and functional state are plausible contributors to response heterogeneity. This agrees with our earlier finding that mitochondrial levels influence apoptotic fate [5], while avoiding the stronger inference that the TMRM maximum itself is MOMP.

We next examined the relation between baseline mitochondrial state and response kinetics. Tmax correlates with the initial TMRM signal, which reflects baseline ΔΨm within an experiment (Figure 1d). Cells with higher initial polarization reach the transition earlier than cells with lower ΔΨm, consistent with the dye reaching the response threshold sooner. The kinetics also depend on light delivery (Figure 1e): increasing pulse frequency at constant intensity per pulse shortens Tmax, following an inverse power-law-like relation. Because each pulse rate was acquired in a separate session, rate is not separable from session and the fitted exponent should be read as descriptive rather than causal.

### An equation parametrises the trajectory of each individual cell

Tmax summarizes the excursion by one number, the position of its maximum, and discards the rest of the curve. To use the whole trajectory we fitted each cell with a model of three free parameters on top of a slow baseline. The measured signal, referenced to the cell’s own starting value, is the sum of a slowly relaxing baseline and a transient that is log-normal in time: a baseline term b + (1 − b)·exp(−t/τ_b), and a transient A·exp(−½·(ln(t/T)/σ)²). The parameters have direct readings. A is the height of the excursion above the baseline, T is the time of its maximum, which is Tmax, and σ sets its shape in logarithmic time. The rise and the decay follow from T and σ: the half-width before the peak is T·(1 − e^(−σ√(2ln2))) and after it T·(e^(σ√(2ln2)) − 1), so the asymmetry of the curve is a property of σ alone (Figure 2a,b).

**Figure 2.**
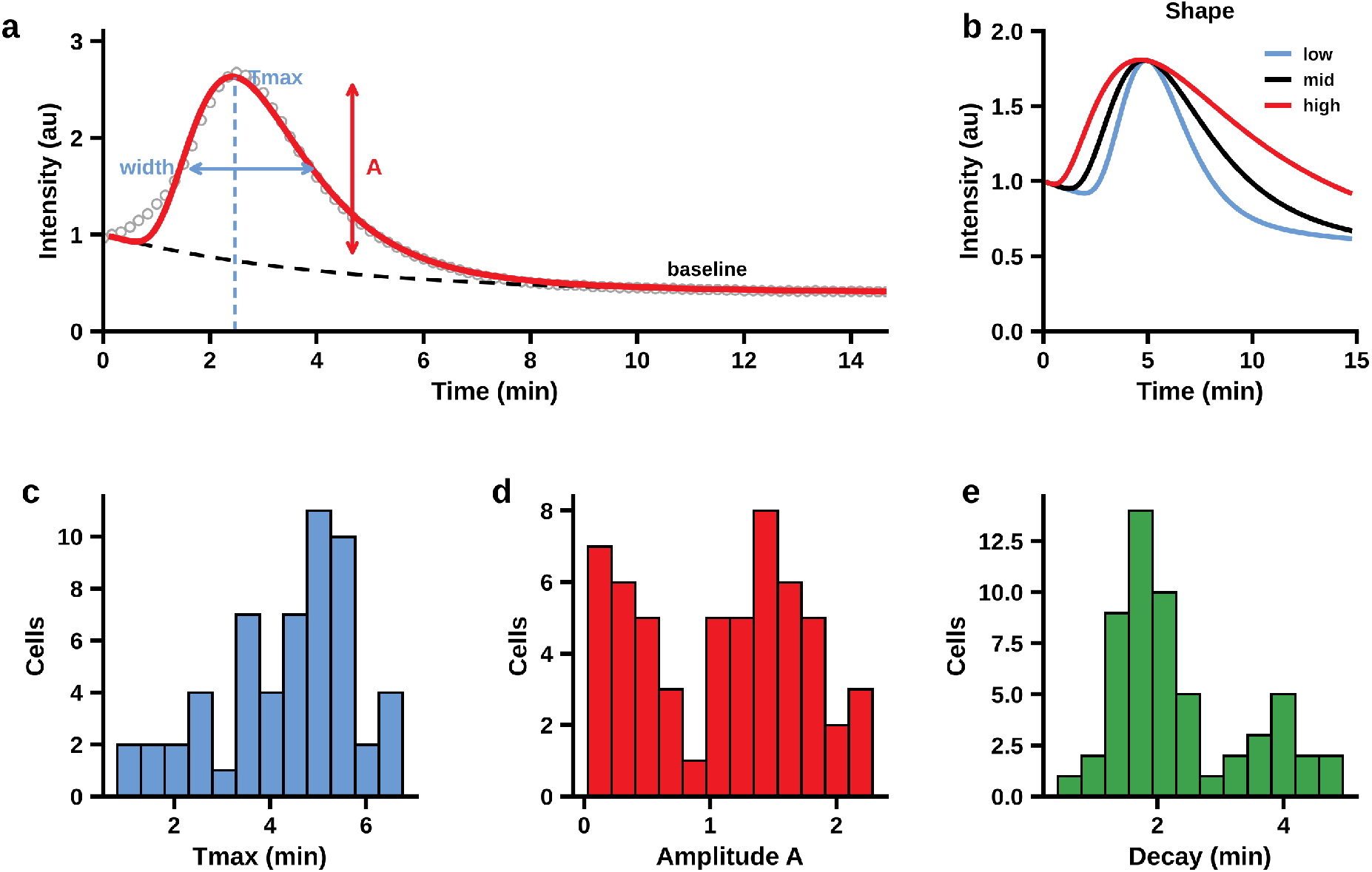
The excursion and its parametrisation. (a) One camptothecin-treated cell. Open circles, measured integrated signal referenced to the cell’s own starting value; red, the fitted total; dashed black, the slow baseline component estimated from the tail of the record and held fixed. Tmax is the position of the maximum, A its height above the baseline and the width the interval between the half-maximum points. (b) Effect of the shape parameter σ on the curve at fixed Tmax and amplitude: larger σ gives a slower rise and a longer decay. (c–e) Distribution across cells of Tmax, of the fitted amplitude and of the decaying half-width.

The baseline and the transient are not separable from a single record. Estimating all five parameters together left the fit poorly conditioned, with b and τ_b correlated above 0.8 in almost every cell. We therefore estimate the baseline from the tail of the record, hold it fixed, and fit only A, T and σ. That reduces the condition number of the problem by nine orders of magnitude while increasing the residual sum of squares by 2.7%, so nothing of consequence is lost. Because the residuals of a trajectory are strongly autocorrelated, uncertainty was obtained by a moving-block bootstrap with the baseline re-estimated inside each replicate, which gives standard errors on T of 11 to 14 s and estimation error small enough that between-cell variation in these parameters is biological rather than instrumental.

Fitting requires enough of the rising phase to be sampled. Records with fewer than about 40 points before the maximum do not constrain σ, and the resulting decay estimates are unreliable. At 10 s intervals this is a requirement on the acquisition rather than on the analysis, and we state it because it governs how such experiments should be designed. Across camptothecin-treated cells the three parameters are distributed as shown in Figure 2c–e: the timing of the excursion is the most uniform property of the population and its amplitude the most variable.

### Which parameter predicts apoptotic fate, and whether any combination does better

Not every cell that showed the excursion went on to die. The photodynamic mitochondrial response is therefore not equivalent to terminal apoptotic commitment. This distinction matters because partial mitochondrial injury can be compatible with survival and, in some settings, can leave genomic damage behind [19,21,22]. Which feature of the trajectory anticipates outcome should therefore be established empirically rather than inferred from the position of the maximum.

We compared seventeen descriptors of the same trajectories against subsequent fate, in 195 cells across four arms: cells that had survived 24 h of camptothecin (59 cells, 43 deaths), two acquisitions of the illuminated no-drug control (41 and 42 cells, 8 deaths each) and TRAIL (53 cells, 24 deaths). The descriptors cover the resting signal, the maximum, the minimum, their difference, the height above the final plateau, the same quantities divided by the resting signal or by cell area, the area of the excursion, the total integral, the time to the maximum, the width of the decay and the initial slope. Because choosing the best of seventeen inflates the apparent result, we built the null distribution of the maximum over the whole search by permutation.

The ranking is consistent and its structure is more informative than its order (Figure 4a). Every descriptor that references the cell’s own resting signal outperforms its raw counterpart: the maximum relative to baseline reaches 0.778 out of sample where the maximum alone reaches 0.688, and the baseline-conditioned response reaches 0.788. The change in emission, the difference between maximum and minimum, predicts fate in its own right (0.751 under camptothecin, 0.735 without drug, 0.758 out of sample), and dividing it by the resting signal improves it slightly. The resting signal by itself carries nothing (0.527), and neither does the total integral (0.515), which is dominated by the baseline. In the camptothecin arm the maximum over the search was 0.779 against a permutation null whose median maximum was 0.637, giving P = 0.008 after correction for the search. The no-drug arm, with eight deaths, does not survive the same correction (P = 0.17), and the TRAIL arm does not either (P = 0.054).

No combination of descriptors improves on the best single one. Fitting penalised logistic models to every pair and every triplet of twelve descriptors, with the standardisation and the coefficients estimated only in the training block and applied without re-estimation, the best single descriptor reached 0.817 out of sample, the best of 66 pairs reached 0.811 and the best of 220 triplets 0.811. The gain from adding a second parameter is −0.006, that is none. This is what the correlation structure predicts and it is worth stating plainly: the trajectory carries one piece of information about fate, not several, so a multivariate score would add complexity without adding discrimination.

The five leading descriptors are rank-correlated between 0.68 and 0.98, so they are one quantity measured five ways. Discrimination does not separate them. Averaged over the three arms imaged with the same protocol they give 0.754 for Fmax / F0, 0.743 for Fmax − Fmin, 0.740 for the height above the plateau, 0.733 for Fmax − F0 and 0.719 for the area of the excursion; resampling the same cells and taking each difference against Fmax − F0 within the replicate, every interval contains zero, the widest running from −0.055 to +0.104. Choosing between them on third decimal places would not survive a second experiment.

A second criterion does separate them, and it decided our choice. The integrated signal of a cell is its mean intensity above background multiplied by its area, and the background is estimated from a region drawn by hand in the same field. When the field is not uniform that estimate carries an error, and the error enters each cell as an additive offset proportional to its area. In a difference the offset appears in both terms and cancels exactly. In a ratio it does not cancel; it biases the ratio towards unity by an amount that depends on the cell. Imposing an inhomogeneous background on our own records confirms the algebra: the area under the curve of every difference is unchanged to three decimal places at background errors up to 10 per cent, while every ratio drifts. Fmax − F0 reached 0.805 out of sample against 0.803 for the ratio, so preferring it costs nothing in discrimination, and it needs no model, no fitting and no cohort-level regression. Two numbers from each trace are enough: where it started and how high it went. The experiment reported below shows how much this matters when the arms being compared were imaged in different fields.

Read as a threshold rather than as a continuous score, the same quantity separates fate cleanly. Taking cells whose signal rose by half above their own resting value as responders, 89.2% of responders died under camptothecin against 45.5% of non-responders, and 30.4% against 5.6% without drug (Figure 3b). This is the single-cell counterpart of the recruitment reported above: priming works by moving cells across this threshold.

**Figure 3.**
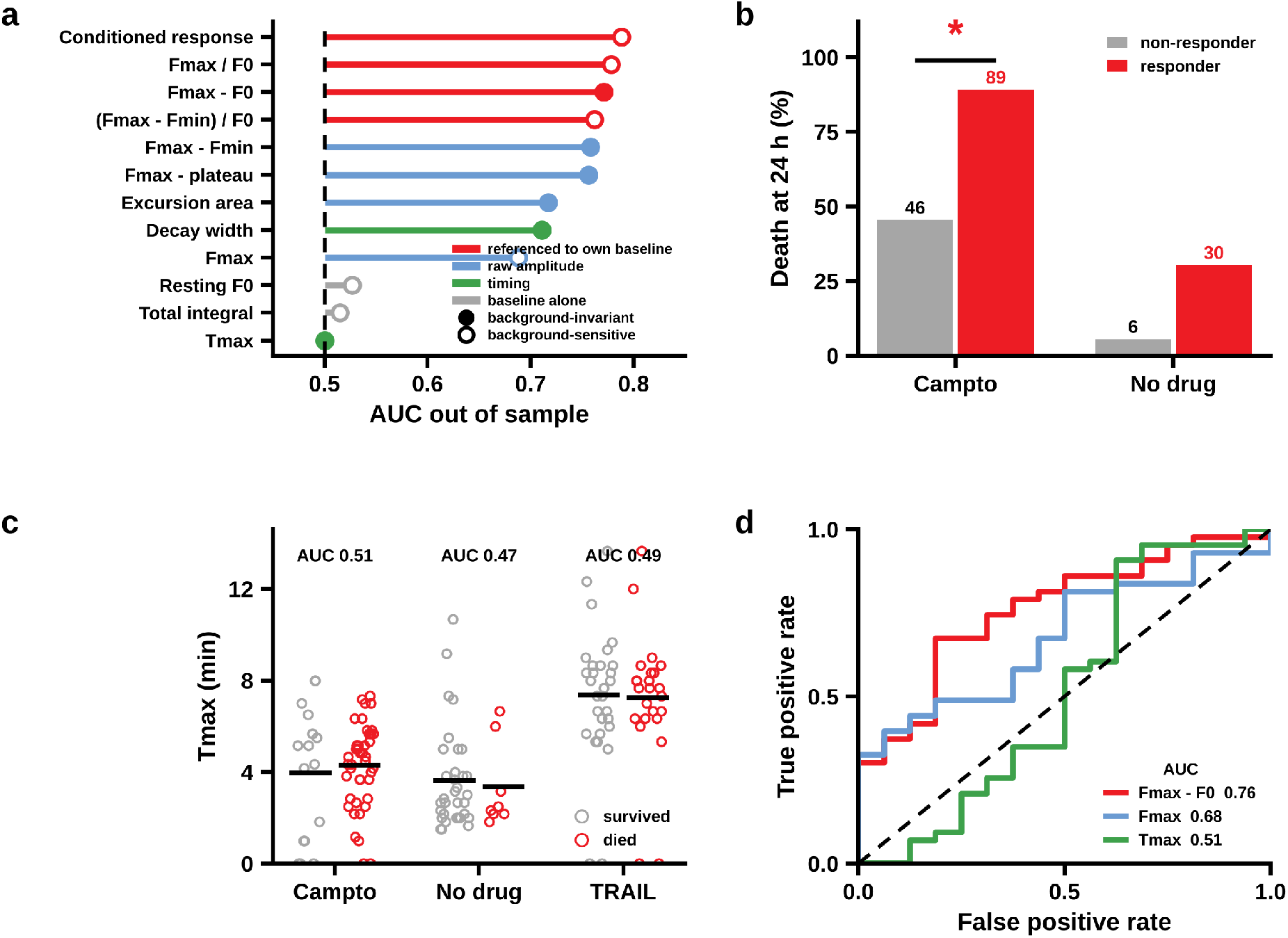
The size of the excursion predicts fate; its timing does not. (a) Seventeen descriptors of the trajectory ranked by their ability to discriminate subsequent death, held out of sample. Red, descriptors referenced to the cell’s own resting signal; blue, raw amplitudes; green, timing; grey, the baseline alone. The dashed line is chance. (b) Death at 24 h among responders and non-responders, where a responder is a cell whose signal rose by at least half above its own resting value, under camptothecin and without drug. (c) Time to the maximum in cells that died and cells that survived, in three arms; the distributions overlap and the areas under the curve are close to chance. Bars are means. (d) ROC curves in the camptothecin arm for the maximum relative to baseline, the maximum alone and the time to the maximum. * P < 0.001.

**Figure 4.**
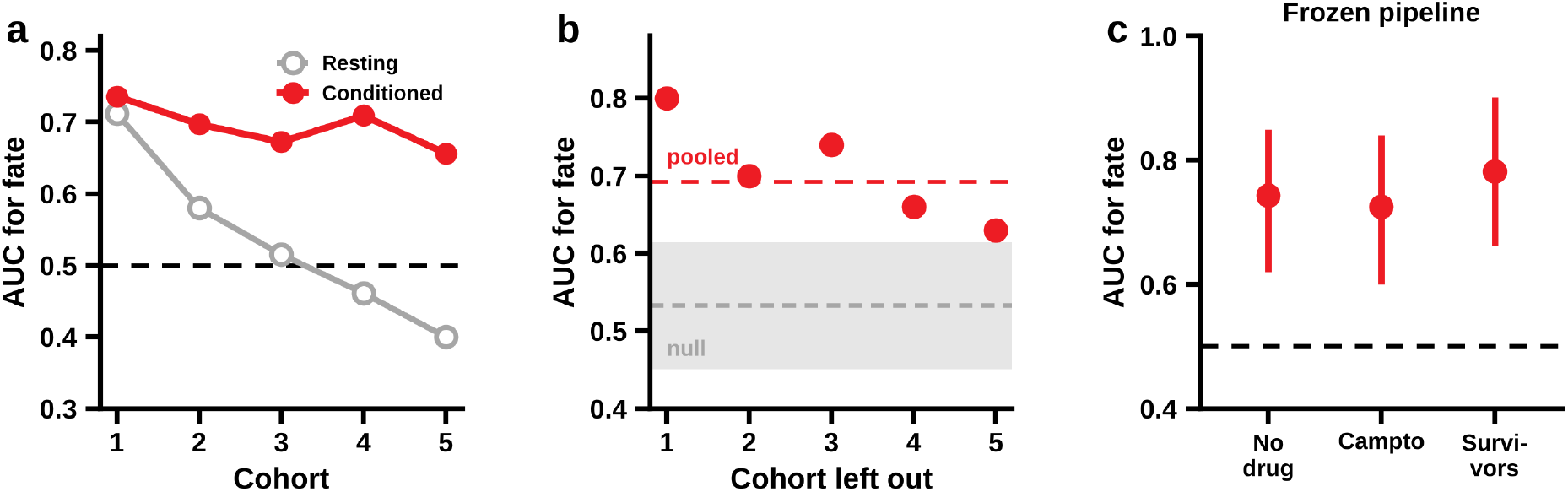
The resting signal does not transfer between experiments; the conditioned response does. (a) Discrimination of subsequent fate in each of five cohorts, 552 camptothecin-treated cells from four cell lines. Grey, resting integrated signal, which crosses 0.5 and reverses direction; red, the baseline-conditioned response, which keeps its direction. (b) Internal–external validation with one cohort withheld, the baseline regression, the standardization and the coefficients estimated only in the remaining cohorts and applied without re-estimation. The dashed red line is the pooled value; the shaded band is the conditional-randomization null, which permutes the response within cohort and within quintile of resting signal, and which no draw of 5,000 reached. (c) A cohort acquired after the analysis was frozen, scored without drug, under camptothecin, and in cells that had already survived 24 h of camptothecin. Bars are 95% confidence intervals.

**Figure 5.**
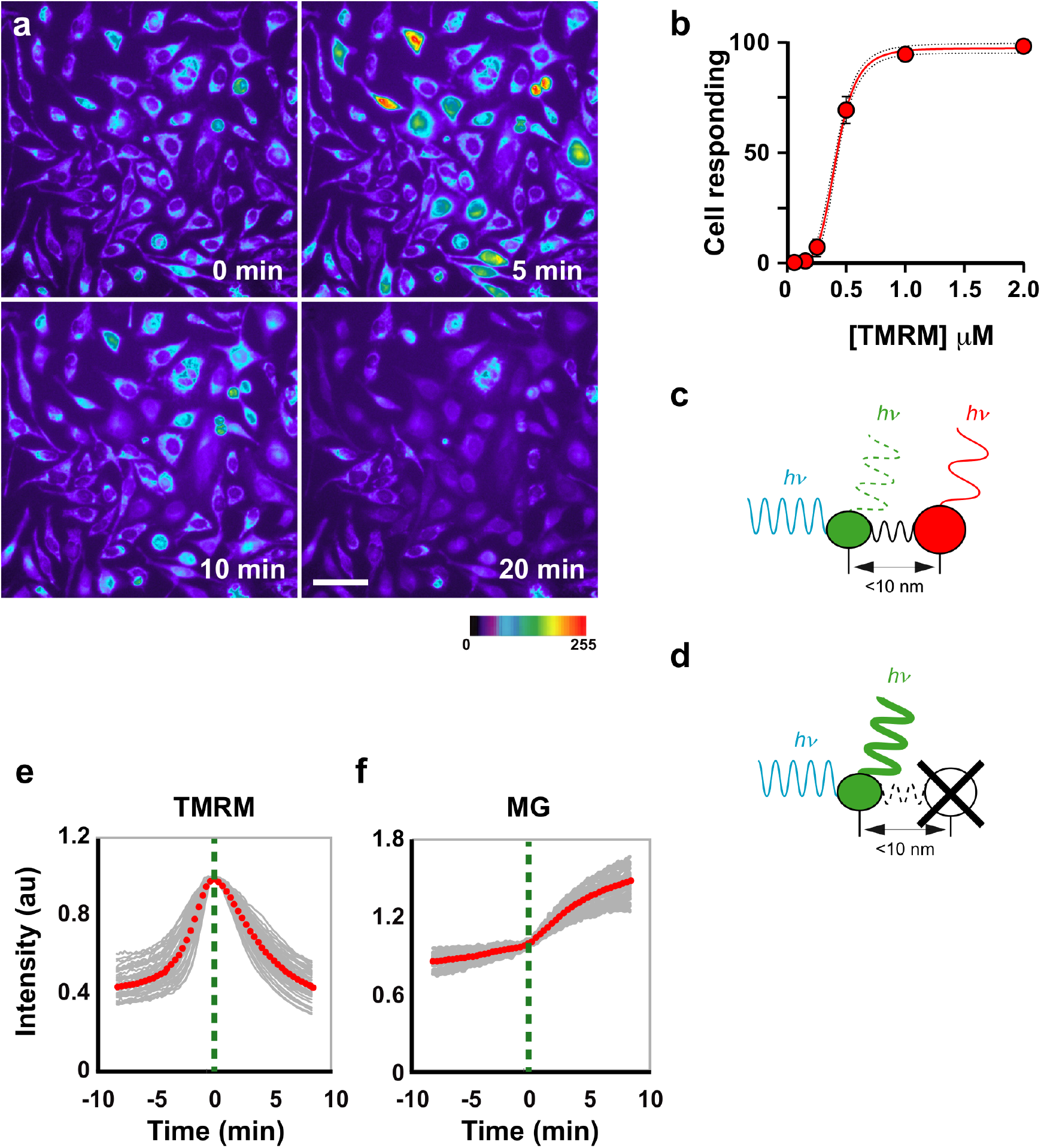
TMRM shows fluorescence enhancement and reciprocal dynamics with MitoTracker Green. HeLa cells were incubated with TMRM for 30 min and filmed at 10 s intervals. (a) Four frames at 0, 5, 10 and 20 min. Some cells undergo a sudden rise in fluorescence followed by decay; one example is marked with red squares. Scale bar 30 µm; fluorescence in pseudocolour. (b) TMRM titration. Cells were incubated with a range of concentrations for 30 min and counted for the excursion. The response follows a Hill curve with a coefficient of approximately 5; dashed lines show the 95% confidence interval. (c,d) Schematic rationale for donor dequenching when an acceptor is lost or photobleached. (e) TMRM time series of 90 cells aligned to their maximum; individual trajectories in grey, mean in red. (f) MitoTracker Green signal for the same cells; red dots show the mean. The MG signal rises near the TMRM maximum, a reciprocal pattern consistent with close fluorophore proximity but not a quantitative demonstration of membrane binding.

The time to the maximum does not discriminate fate in any of the four arms, with areas under the curve of 0.511, 0.468, 0.461 and 0.486, and the distributions of Tmax for cells that died and cells that survived overlap almost completely (Figure 3c,d). We report this because an earlier acquisition from this laboratory, with records long enough for Tmax to spread across half an hour, had suggested a threshold near ten minutes. In the records analysed here the median Tmax is 2.7 to 7.7 min and more than 97% of cells fall on one side of that threshold, so it separates almost nothing. Whether timing carries information when it is given room to vary is a question these data cannot answer, and we make no claim for it.

### The predictor transfers between experiments; the resting signal does not

Figure 1d shows that Tmax and the initial TMRM signal are related within one experiment. That relation raises a question for any use of the resting signal as a biomarker, because dye loading depends on concentration, on loading time, which in our hands ranges from 30 to 60 min depending on the day and the operator, and on the excitation intensity. We therefore asked whether the resting signal predicts fate across experiments, or only within one.

In 552 camptothecin-treated cells from four cell lines, acquired in five separate cohorts, the resting integrated signal discriminated subsequent death with areas under the curve ranging from 0.40 to 0.82, and it reversed direction between cohorts, crossing 0.5 (Figure 4a). A predictor that changes sign between experiments cannot be used in a new one. The excursion conditioned on the cell’s own resting signal behaves differently. Taking the standardized residual of the logarithm of the peak on the logarithm of the baseline, fitted within each cohort, gives the same direction in all five, with areas under the curve from 0.656 to 0.736 and a pooled value of 0.689.

Because the residual is fitted within cohort, it must be validated with a cohort held out entirely. Estimating the baseline regression, the standardization and the coefficients only in the remaining cohorts, and applying them without re-estimation, gives 0.63 to 0.80 across the five held-out cohorts (Figure 4b). To exclude the possibility that this reflects residual information in the baseline, we permuted the residual within cohort and within quintile of resting signal: none of 5,000 randomizations reached the observed value (P = 2 × 10⁻⁴), and the shaded band in Figure 9b shows that null. A cohort acquired after the analysis was frozen reached 0.743 scored without drug and 0.725 under camptothecin, and in cells that had already survived 24 h of camptothecin the conditioned response gave 0.782, with a hazard ratio of 1.48 per standard deviation for the timing of death (Figure 4c).

Two consequences follow for the use of this measurement. The resting intensity of a mitochondrial dye should not be carried between experiments as a predictor, whatever it achieves within one. And conditioning the response on the cell’s own starting point removes most of what differs between days, which is what allows a threshold fitted in one experiment to be applied in the next.

### The excursion is accompanied by ROS production

ROS can contribute to mitochondrial depolarization and apoptotic commitment [20], so we asked whether TMRM generates them under controlled irradiation. We followed ROS in real time with CellROX Green, which emits more strongly on oxidation. CellROX fluorescence rose in parallel with TMRM fluorescence (Figure 6a,b), both with sigmoidal kinetics: a lag, rapid growth and then a plateau.

**Figure 6.**
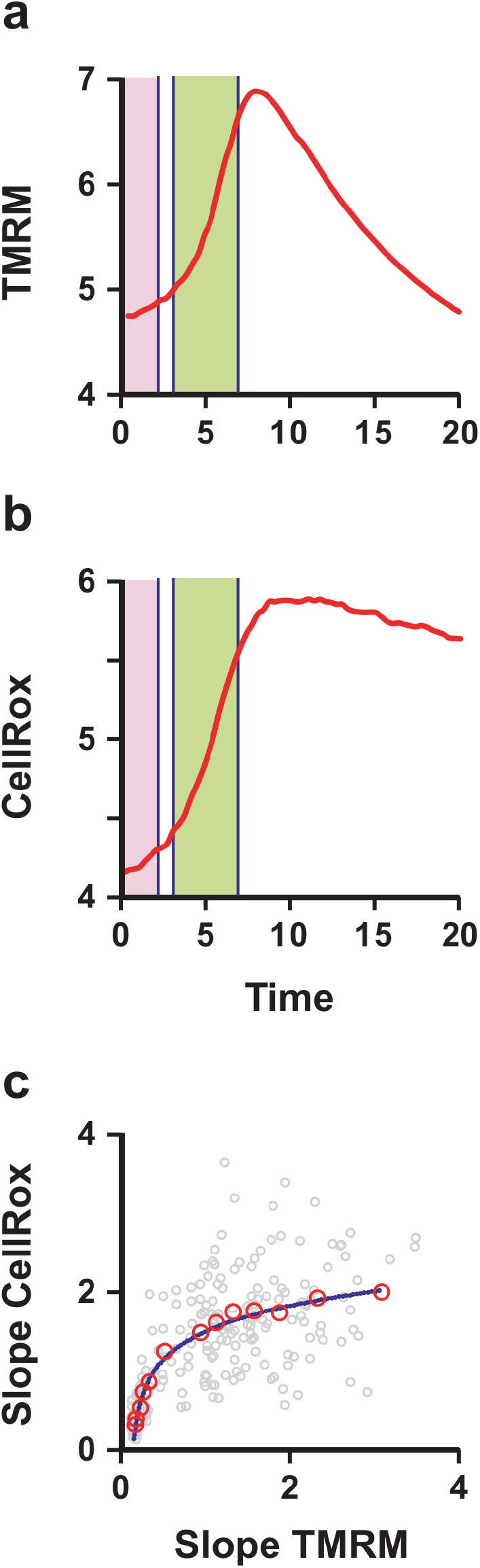
TMRM fluorescence enhancement is associated with ROS production. Cells were co-incubated with TMRM and CellROX. (a) TMRM trajectory in a single cell. (b) CellROX signal in the same cell. Both rise in parallel during the rising phase of TMRM (green zone), which allows the two growth rates to be compared cell by cell. (c) The two slopes plotted against the TMRM slope, showing a logarithmic relation. Individual cells in grey; means obtained by stratified sampling in red, with a black trend line.

Comparing the rising phases cell by cell, the slope of TMRM relates to the slope of CellROX logarithmically (Figure 6c). ROS production rises in proportion to TMRM activity at low activity and then increases much more slowly at higher activity. The saturation is what one would expect from local oxygen depletion or from self-quenching of the dye, either of which would limit ROS output while the photosensitizer continues to absorb light. Two conclusions follow. ROS generation is driven directly by TMRM excitation in the linear regime, and above a threshold the yield is limited by the microenvironment rather than by the dye. Irradiation protocols for therapy should therefore be set to avoid that saturation.

### Glycolysis and PKM2 set how many cells enter the responding state

Mitochondria control both metabolism and cell death, and the two are linked at several levels; imbalances in the acetyl-CoA/CoA and NAD+/NADH ratios trigger signalling that determines fate [23]. To find the origin of the variability in response within a clonal population, we looked at energy metabolism. ΔΨm reports mitochondrial function, and its variability within a clone plausibly reflects metabolic heterogeneity. The Krebs cycle sustains ΔΨm by generating NADH and FADH₂, which donate electrons to the electron transport chain and power proton extrusion. Glycolysis is cytosolic, but its end product, pyruvate, enters mitochondria through the mitochondrial pyruvate carrier and is converted by pyruvate dehydrogenase to acetyl-CoA, which fuels the cycle. Glycolytic activity therefore has a direct route to ΔΨm.

If that route is real, raising intracellular pyruvate should raise ΔΨm in the cells that start low, and should therefore carry some of them above the threshold at which the dye reaches the concentration at which the excursion occurs. The prediction is about how many cells respond, not about how fast they respond. We tested it with ethyl pyruvate (EP), a membrane-permeable analogue that raises intracellular pyruvate [24], and with TEPP-46, an activator of PKM2 [25].

Both treatments increased the fraction of cells showing photo-enhancement (Figure 7a,b). EP at 10 mM for 5 h raised the responding fraction above the untreated control, and TEPP-46 at 10 µM for 5 h raised it further, until almost every cell in the field responded. Because the non-responding cells are enriched among those with lower initial ΔΨm (Figure 1b), the additional responders are consistent with cells being recruited across the response threshold rather than only with changes in cells that already responded. Tmax also fell by about 35% under both treatments (Figure 7c), and the spread of Tmax narrowed (Figure 7d). These latter observations are not sufficient to establish variance control as an independent effect, but they raise the possibility that metabolic priming can constrain cell-to-cell response variability as well as shift the responding fraction.

**Figure 7.**
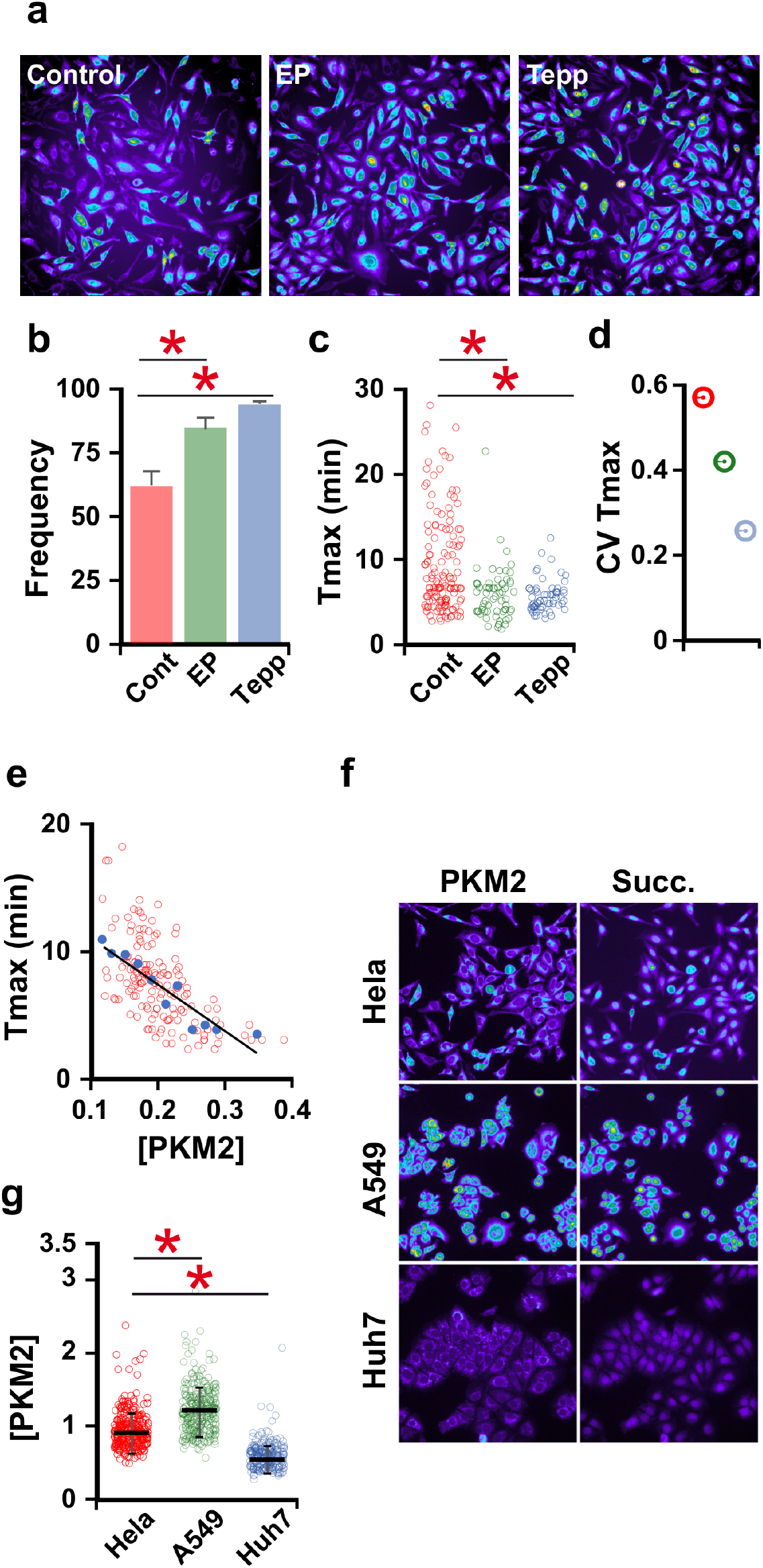
Metabolic state modulates entry into the TMRM-responsive state. (a) Maximum-intensity map of control HeLa cells over the experiment; cells of high intensity are those that showed the excursion. EP, 10 mM ethyl pyruvate for 5 h; Tepp, 10 µM TEPP-46 for 5 h. (b) Percentage of cells showing the excursion under each condition; both treatments increase the responding fraction, and under TEPP-46 almost the whole population responds. (c) Distribution of Tmax across individual cells. (d) Coefficient of variation of Tmax; narrowing under TEPP-46 is reported as an observation rather than as an independently established variance-control effect. (e) Tmax against PKM2 concentration measured after fixation and immunofluorescence in the same re-identified cells. Higher PKM2 is associated with lower Tmax. (f) PKM2 concentration, normalized to cell volume, across cell lines. (g) Relative PKM2 concentration in individual cells. * P < 0.001.

Recruitment has a direct consequence for the damage delivered. A responding cell is one in which the dye has reached that concentration, and it is there that illumination produces ROS inside its mitochondria (Figure 6). Raising the responding fraction therefore raises the number of cells in the population that sustain mitochondrial oxidative damage during irradiation. The mitochondrial compartment is unusually exposed to that damage, because its proteins are statistically depleted in reactive cysteine and so carry less of the buffering that protects other compartments [43]. This is the simplest explanation for why priming improves the outcome of the combination treatments (Figure 9c, groups 5 and 6): a larger share of the population enters chemotherapy already carrying mitochondrial damage, and therefore with a lowered apoptotic threshold. We note that the experiments measure recruitment and killing in separate populations and do not follow the same cells through both, so this remains an inference rather than a demonstrated chain.

To relate the response to PKM2 at single-cell resolution, we photoactivated TMRM, then fixed the cells and stained for pyruvate kinase M2 (PKM2), the enzyme that catalyses the final step of glycolysis. PKM2 abundance correlates negatively with Tmax (Figure 7e). Cells with high PKM2 reached peak fluorescence two to three times faster than PKM2-low cells (r = −0.89, P < 0.001). This establishes an association between PKM2 abundance and response kinetics in the same cells; it is consistent with, but does not by itself prove, a causal chain from glycolytic pyruvate production to ΔΨm and photodynamic response. Across cell lines, A549, in which approximately 90% of cells respond, has higher PKM2 concentration, whereas Huh7, in which only approximately 2% respond, has much lower PKM2 (Figure 7f,g).

Metabolic preconditioning through PKM2 activation could therefore alter both the fraction of cells entering the response and, potentially, its heterogeneity. To place PKM2 in clinical context we examined its prognostic weight across malignancies with the GEPIA platform [26] (Figure S1). A meta-analysis of 27 studies and 4,796 patients with solid tumours reports that high PKM2 is associated with reduced overall survival (HR 1.73, 95% CI 1.48–2.03) and disease-free survival (HR 1.90, 95% CI 1.39–2.59) [27], and the association has also been reported in multivariable analyses [28]. In glioblastoma and lung cancer, high PKM2 expression is likewise associated with poorer outcomes [29,30]. These clinical associations do not demonstrate susceptibility to the present intervention, but they support PKM2 as a biologically relevant axis on which to test metabolic recruitment.

Given the spread of PKM2 across cell lines, we examined its heterogeneity in clinical material. Quantitative immunofluorescence was performed on paraffin sections of human skin squamous cell carcinoma (SCSC) and basal cell carcinoma (BCC), normalizing PKM2 to cell mass and measuring it cell by cell (Figure 8a,b). SCSC shows substantial intratumoural variability in PKM2, whereas BCC shows markedly less (Figure 8c). These data do not establish PKM2 as a clinical stratification biomarker, but they show that the relevant heterogeneity is measurable in tumour tissue and motivate prospective evaluation of PKM2 level and distribution in future translational studies.

**Figure 8.**
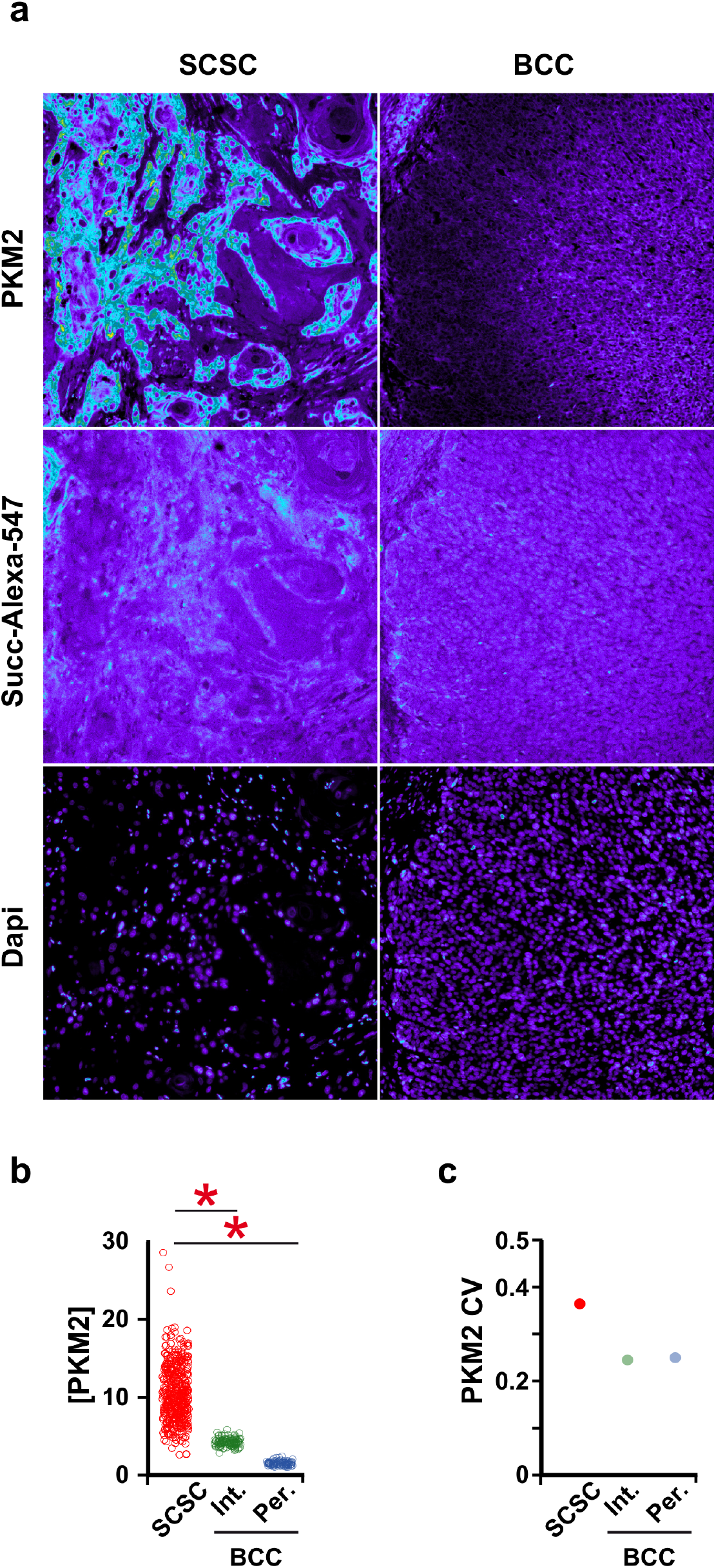
PKM2 is heterogeneously expressed in human tumours. (a) PKM2 in squamous cell carcinoma of the skin (SCSC, left) and basal cell carcinoma (BCC, right), in pseudocolour. Middle row, succinimidyl ester–Alexa 547, which labels protein amino groups and provides a reference for total protein. Bottom row, DAPI. (b) PKM2 concentration in SCSC tumour cells (red), in the inner region of BCC (green) and at the BCC periphery (blue). (c) Coefficient of variation of PKM2 across the tumour types.

**Figure 9.**
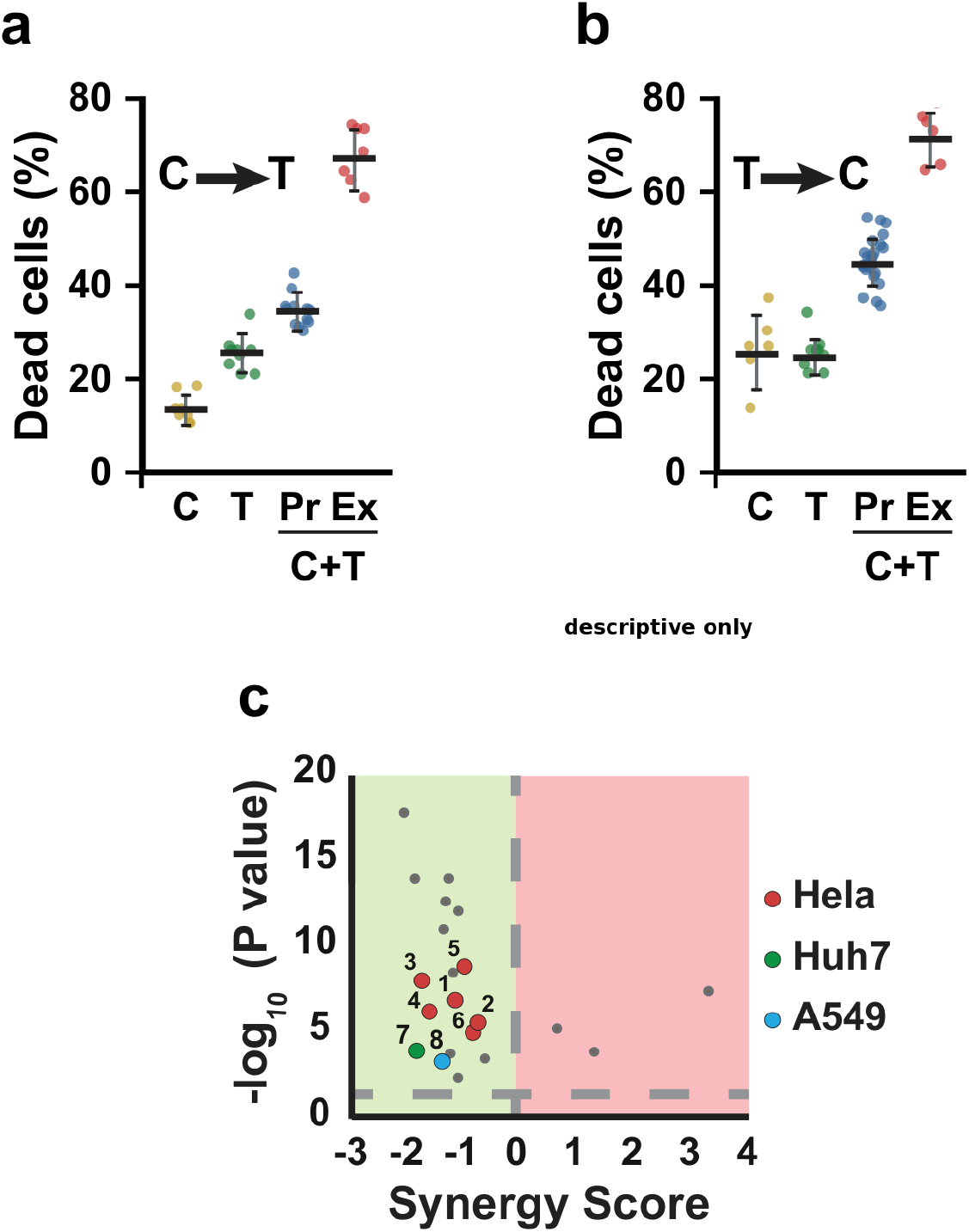
TMRM photodynamic treatment increases killing in combination with camptothecin or TRAIL. (a) Cells first exposed to 2.5 µM camptothecin for 24 h and then treated with TMRM-PDT (C to T). Yellow, death from camptothecin alone; green, death from TMRM-PDT alone; blue, Bliss-independence expectation; red, observed combination death. (b) Reverse order: TMRM-PDT first, then 2.5 µM camptothecin for 24 h (T to C). (c) Bliss-deviation scores for combinations and cell lines. Red, HeLa; green, Huh7; blue, A549. Grey dots are reference values published previously by our group [4]. Groups: 1, HeLa, camptothecin then TMRM-PDT; 2, HeLa, TMRM-PDT then camptothecin; 3, HeLa, TMRM-PDT then TRAIL; 4, HeLa, TRAIL then TMRM-PDT; 5, HeLa, TEPP-46 then TMRM-PDT; 6, HeLa, ethyl pyruvate then TMRM-PDT; 7, Huh7, TMRM-PDT then camptothecin; 8, A549, TMRM-PDT then camptothecin. All tested combinations fell below the Bliss-independence expectation in their corresponding experiment. Because each arm was measured once, these values are descriptive and no inferential P value is assigned to synergy.

### Chemotherapy moves cells into the responding state and identifies a candidate intervention window

If the excursion reports a state rather than a fixed property, the state should change during treatment, and there should be a moment at which most of the population is in it. That moment is what would decide when to apply the optical challenge. We tested it by pre-incubating cells with 5 µM camptothecin for 4 h and then loading TMRM, against an untreated control acquired in the same session and scored with the same background region.

Four hours of camptothecin shifts the population (Figure 10a,b). The median of Fmax − F0 rises by a factor of 1.498 (95% interval 1.23 to 1.66), and the proportion of cells above the control median rises from 50% to 76% (difference 26.3 percentage points, 15.8 to 36.3). The same contrast measured with the other background-invariant descriptors is smaller: 70% for Fmax − Fmin, 70% for the area of the excursion, 68% for the height above the plateau. Segmented cell area is unchanged between the arms, so this is not a difference in what was measured. The effect is in the direction the earlier results predict: a treatment that hyperpolarises mitochondria should carry cells across the threshold at which the dye concentrates enough to respond, and it does.

**Figure 10.**
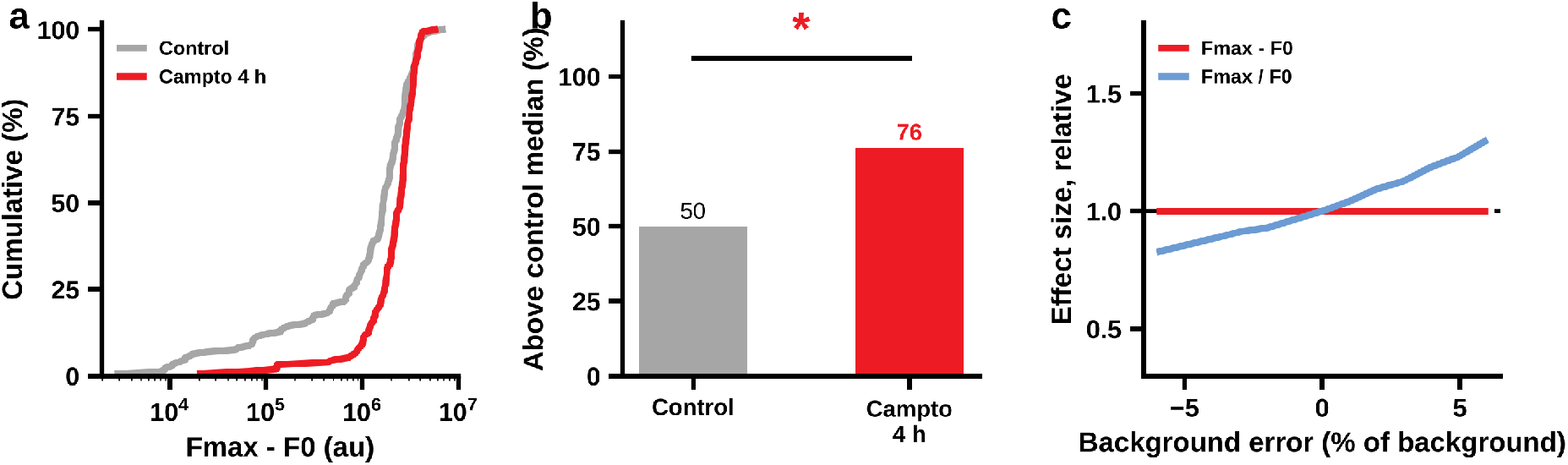
Chemotherapy moves cells into the responding state. HeLa cells pre-incubated with 5 µM camptothecin for 4 h and then loaded with TMRM, against an untreated control acquired in the same session; 172 and 152 unique trajectories after removing duplicated columns from the export. (a) Cumulative distribution of Fmax − F0 in the two arms. (b) Proportion of cells above the median of the control. (c) Robustness to an error in background estimation: the effect size measured as a difference is invariant, the same effect measured as a ratio is not. * P < 0.001.

The experiment also shows why the difference rather than the ratio should be used. The two arms were imaged in different fields, and their median background intensities differ by 28 per cent, so there is no reason for the two background estimates to carry the same error. Introducing a systematic error in one arm leaves the ratio of Fmax − F0 between arms at 1.498 at every level tested, from one per cent to ten. Over the same range the ratio of Fmax / F0 moves from 1.78 to 1.92 at one per cent, from 1.63 to 2.19 at four, and from 1.34 to 2.83 at ten (Figure 10c). Measured as a ratio, then, the apparent size of this effect is set as much by how carefully the background polygon was drawn as by the biology, and in a smaller experiment its sign would be too. That is the whole argument for the difference. Among the descriptors that are invariant to the background, Fmax − F0 also gives the largest separation between the arms: 1.498 against 1.291 for the area of the excursion, 1.276 for Fmax − Fmin and 1.229 for the height above the plateau, and each of those three differences from Fmax − F0 is itself outside the resampled interval. The quantity that is hardest to corrupt is also, here, the most sensitive one.

Two limitations bear on how far this can be taken. The resting signal is lower in the treated arm by a factor of 0.60, which is the opposite of what mitochondrial hyperpolarisation alone would produce and which we cannot currently explain; it may reflect differences in dye loading between wells, or depolarisation by four hours at this concentration, and it is one of the reasons we do not build on the resting signal. And this is a single pair of arms at a single delay. Establishing the window as a curve, rather than as one point on it, requires the same comparison at several delays within each session, which is the experiment we would do next.

Taken with the order effect reported below, in which the optical challenge was more effective when applied before camptothecin than after a full day of exposure, the result supports a practical hypothesis: the useful quantity to follow through treatment may be the fraction of cells whose excursion exceeds a defined response threshold rather than the resting intensity itself. This experiment samples only one delay, so it identifies a candidate window rather than defining the full temporal window. A true timing rule will require within-session measurements across several delays.

### Combination treatment increases killing and shows a timing dependence

Finally, we asked whether TMRM photodynamic treatment increases killing when combined with chemotherapy. In one experiment, cells were exposed to camptothecin, loaded with TMRM 24 h later, photoactivated and then followed by time-lapse phase-contrast microscopy. Combination survival fell below the value expected under Bliss independence and below survival with either treatment alone (Figure 9a,c). Because each arm was measured once in this experiment, the Bliss comparison is descriptive rather than evidence of reproducible synergy.

In a second set of experiments, cells were loaded with TMRM and irradiated before camptothecin or TRAIL was added. This order also increased killing, and the observed combination survival again fell below the Bliss-independence expectation (Figure 9b,c). The pattern was seen in three cell lines, including Huh7, in which the characteristic excursion is rarely observed.

These observations show that enhancement of killing does not depend strictly on Tmax. A plausible explanation is that photodynamic mitochondrial injury lowers the apoptotic threshold and increases subsequent drug sensitivity, consistent with the general concept of mitochondrial priming [38]. The Huh7 result also cautions against equating the visible excursion with the whole biological effect of TMRM photoactivation. Independent biological replication will be required to determine the magnitude and reproducibility of the combination effect and whether it meets a formal criterion for synergy.

## Discussion

Treatment resistance persists because tumour populations are heterogeneous and plastic. Previous work established that TMRM can function as a mitochondria-targeted photosensitizer [46]. The contribution of the present study is different: we identify a dynamic, perturbation-revealed mitochondrial state that varies between cells, predicts subsequent fate better than resting intensity, and can be pharmacologically recruited by metabolic priming.

The response is favoured in cells with higher mitochondrial polarization, where TMRM accumulates more strongly. Illumination then produces ROS and mitochondrial injury. The resulting selectivity is therefore conditional on both cell state and light delivery rather than being an intrinsic tumour-targeting property of the dye. This distinction is important because it shifts the emphasis from TMRM as a photosensitizer per se to the metabolic state that determines whether a cell enters the vulnerable regime.

The most transferable information is not the resting TMRM signal. Resting intensity varies substantially between experimental cohorts and even reverses its relation to fate, whereas the response conditioned on the cell’s own starting point preserves its direction. How far the excursion rises above baseline predicts subsequent death; when it peaks does not in the datasets analysed here. Thus the relevant phenotype is revealed by perturbation rather than reliably encoded in a static measurement. Fmax − F0 is attractive not because it wins by a marginal difference in AUC, but because an additive background error cancels in the difference and the measurement can therefore be carried more safely between fields and experiments.

The reciprocal dynamics of TMRM and MitoTracker Green are consistent with close proximity of the two fluorophores in the mitochondrial membrane environment, but they should not be read as a definitive demonstration of TMRM binding to the inner membrane. The mechanistic conclusion required for the present study is narrower: TMRM accumulation is voltage dependent, repeated illumination produces a local mitochondrial response, and ROS rise during that response. The principal biological claims do not depend on assigning TMRM to a specific membrane-binding mode.

We likewise make no claim about the precise photochemical route of killing. Rhodamines can generate radicals through electron or hydrogen transfer and can also transfer energy to molecular oxygen. Saturation of ROS output at high TMRM activity is compatible with local oxygen limitation or self-quenching, among other possibilities. Experiments in deuterated medium and with singlet-oxygen quenchers could distinguish these mechanisms, but they are not required for the central result: the dynamic response predicts fate, metabolic priming recruits cells into the responding state, and combination treatment increases killing beyond that observed with either treatment alone in the experiments performed.

In the combination experiments, survival fell below the Bliss-independence expectation for camptothecin and TRAIL and in both treatment orders. Because these measurements were single determinations per arm, we interpret them as a consistent pattern across the tested conditions rather than as a statistically established synergy claim. A plausible mechanism is that photodynamic mitochondrial injury lowers the apoptotic threshold, consistent with mitochondrial priming [38], but direct demonstration of that causal chain will require experiments that follow recruitment, mitochondrial injury and fate in the same cells.

Metabolic state controls how many cells enter the responding regime. TEPP-46 and ethyl pyruvate increase the responding fraction, and PKM2 abundance is associated with faster response kinetics in the same cells. These observations are consistent with a model in which glycolytic pyruvate supply modulates mitochondrial state and thereby the probability of crossing the photodynamic response threshold. They do not establish PKM2 as the sole causal determinant. Of particular interest, TEPP-46 also narrows the Tmax distribution. The present experiments were not designed to test variance control as an independent endpoint, but this observation raises a broader possibility: metabolic priming may alter not only the population mean or responding fraction, but also the dispersion of single-cell responses.

High PKM2 expression is associated with poor outcome in several tumour types [29,41,42], and our tissue measurements show that PKM2 can be highly heterogeneous within human tumours. These findings make PKM2 a candidate marker for future stratification studies, but they do not establish that PKM2-high tumours will be preferentially susceptible to TMRM-PDT. That prediction should be tested prospectively with clinically relevant photosensitizers and patient-derived models.

Which quantity to measure deserves a paragraph of its own, because the choice is not settled by discrimination. Five descriptors of the excursion predict fate equally well, within a resampled interval that comfortably contains zero for every pairwise difference, and they are rank-correlated between 0.68 and 0.98; they are one quantity seen five ways. What separates them is how they behave when the background is not estimated perfectly. Any integrated single-cell signal is a mean intensity above background times an area, so an error in the background enters as an offset proportional to cell area: present in both terms of a difference and absent from it, present in only the numerator and the denominator of a ratio and therefore not removed. In our own 4 h experiment, where the two fields differ in background by 28 per cent, the effect measured as Fmax − F0 is 1.498 at every simulated error level while the same effect measured as Fmax / F0 ranges from 1.34 to 2.83. A quantity whose value depends on how a background polygon was drawn is not one to carry into a clinical measurement, however well it performs on a clean dataset. We therefore recommend Fmax − F0, and we would make the same recommendation for any assay that integrates intensity over a segmented cell.

The selectivity of this approach is worth stating precisely. It is not conventional pharmacological targeting: TMRM itself has no tumour-specific ligand. Selectivity arises conditionally from the intersection of where light is delivered and which cells have accumulated enough dye to enter the vulnerable response. This makes the intervention spatially and temporally controllable, as in established photodynamic therapy. Light penetration therefore restricts direct application to accessible sites such as skin, bladder, oesophagus and endoscopic or intraoperative fields, and TMRM itself is not a clinical agent. Translation would require clinically suitable mitochondria-targeted photosensitizers. These limitations do not affect the mechanistic conclusion that metabolic state can determine entry into a photodynamically vulnerable regime.

The main limitation is the absence of in vivo validation. The work is a mechanistic and methodological proof of concept centred on single-cell heterogeneity and dynamic vulnerability. The pulse-rate series was acquired with one rate per session, so the fitted scaling exponent is descriptive and does not separate pulse rate from session. The combination experiments contain single determinations per arm; Bliss deviations should therefore be treated as descriptive and independently replicated before claiming reproducible synergy. The reciprocal TMRM–MitoTracker Green signal is consistent with energy transfer but does not exclude other explanations, and the TMRM maximum should not be equated with MOMP without an independent permeabilization marker. Finally, the 4 h camptothecin experiment samples one delay only, so it identifies a candidate intervention window rather than the full temporal trajectory. These limitations define the next experiments rather than altering the central observation that a dynamic mitochondrial vulnerability can be measured and pharmacologically recruited.

In summary, TMRM photoactivation exposes a dynamic mitochondrial vulnerability whose magnitude relative to each cell’s own baseline predicts subsequent fate. Static resting intensity does not transfer reliably between experiments, whereas the perturbation-evoked response does. Metabolic priming with TEPP-46 or ethyl pyruvate recruits otherwise non-responding cells across the response threshold, and chemotherapy itself shifts the population distribution over time. Together, these results support a general principle: a therapeutically relevant state can be revealed by perturbation, pharmacologically recruited and potentially exploited during a transient window before resistant populations become stabilised [47].

## Materials and methods

### Cell lines and materials

HeLa (ATCC CCL-2), A549 (ATCC CCL-185) and Huh7 (CVCL_0336) cells were grown in Dulbecco’s Modified Eagle Medium (DMEM, Gibco) GlutaMAX-I supplemented with 10% foetal bovine serum (FBS, HyClone) and penicillin–streptomycin (Sigma), at 37 °C in a humidified incubator with ∼5% CO₂. For particular experiments DMEM without glutamine or without FBS was used. Treatments: TEPP-46 (Sigma-Aldrich #505487) and ethyl pyruvate (Sigma-Aldrich #E47808).

### TMRM staining and photoactivation

Mitochondria were stained with TMRM (Molecular Probes T668) at the concentrations given in the figures, for 30 min at 37 °C. Cells were then washed three times with pre-warmed culture medium and transferred immediately to an inverted fluorescence microscope. For photoactivation, cells were irradiated with a CoolLED source at 50% power through a rhodamine filter set (excitation 548 nm, emission 573 nm). Time-lapse images were acquired at 10 s intervals for 30 min at 20× magnification (0.4 NA HCX PL FL) on a widefield Leica DMi8 (Leica Microsystems) with a Hamamatsu ORCA-Flash4.0 LT sCMOS camera. Integrated TMRM fluorescence per cell was quantified with MetaMorph 7.8.0.0 (Molecular Devices). Background was subtracted using adjacent cell-free regions and the data were normalized to the baseline measured before photoactivation.

### Reciprocal TMRM–MitoTracker Green imaging during photoactivation

Cells were co-stained with MitoTracker Green (Molecular Probes) at 10 ng/mL and TMRM at 500 nM, applied together in culture medium for 30 min at 37 °C under 5% CO₂, then washed three times with pre-warmed medium. Imaging used 488 nm excitation with 525/50 nm emission for MitoTracker Green and 561 nm excitation with 605/70 nm emission for TMRM, on a temperature-controlled stage at 37 °C, every 10 s for 30 min with an ORCA-Flash4.0 LT sCMOS camera. Integrated fluorescence per cell was quantified in MetaMorph, with background subtraction from adjacent cell-free regions and normalization to baseline. The experiment tests for reciprocal donor/acceptor-like dynamics during TMRM photoactivation; because no independently calibrated FRET efficiency was obtained, these measurements are interpreted as evidence consistent with close fluorophore proximity rather than as a quantitative binding assay.

### Live-cell microscopy and fate tracking

HeLa cells were seeded in 24-well plates (Nunc) the day before the experiment and imaged every 30 min for 24 h at 37 °C in a humidified chamber with ∼5% CO₂, at 20× magnification (0.4 NA HCX PL FL) on a Leica DMi8 with an ORCA-Flash4.0 LT camera. Images were analysed in MetaMorph 7.8.0.0. Time to death was scored from the morphological changes of apoptosis. Apoptosis was triggered with human recombinant TRAIL (Millipore) at 7.5 ng/mL or camptothecin at 2.5 µM.

### Cell volume

Cell volume was estimated from total protein content as described [43,44]. Total protein was labelled with succinimidyl ester coupled to Alexa 647 (Invitrogen), which forms covalent bonds with primary amines. After immunolabelling, cells were incubated with 0.2 µg/mL of the ester in PBS for 5 min, washed and mounted. The signal from an individual cell is proportional to its volume [43,44].

### Immunostaining and wide-confocal cytometry

Cells growing on coverslips (HeLa, Huh7, A549) were fixed and PKM2 was labelled indirectly with a mouse anti-PKM2 primary antibody (Santa Cruz, C-11, sc-365684) and an Alexa Fluor 488 donkey anti-mouse IgG (H+L) secondary (Invitrogen). Coverslips were mounted in Vectashield (Vector Laboratories). Images were collected on a Leica TCS SP5 multispectral confocal system with a 20× 0.7 HCX PL APO CS objective and the pinhole fully open, to collect the maximum light emitted by the specimen. Hundreds of cells across different fields were collected per slide and analysed in MetaMorph 7.8.0.0.

For tumour immunohistochemistry, biopsies were formalin-fixed and paraffin-embedded. Sections of 5 µm were treated with EnVision FLEX Target Retrieval Solution, low pH (DAKO), at 95 °C for 2 min to unmask antigens, and then labelled as for cultured cells.

### Trajectory model and its estimation

Each per-cell trace was referenced to the median of its first three points. The model is y(t) = b + (1 − b)·exp(−t/τ_b) + A·exp(−½·(ln(t/T)/σ)²). The baseline parameters b and τ_b were estimated by least squares on the last 30% of the record and then held fixed; A, T and σ were estimated on the whole record by Levenberg–Marquardt from several starting points, retaining the best fit. Derived quantities are the rising half-width T·(1 − e^(−σ√(2ln2))), the decaying half-width T·(e^(σ√(2ln2)) − 1) and their sum. Uncertainty was obtained by a non-circular moving-block bootstrap of the residuals with blocks of 30 points and the baseline re-estimated within each replicate. Cells were retained when the maximum fell inside the record, when at least 40 points preceded it and when the residual root mean square was small relative to the fitted amplitude.

### Baseline-conditioned response and its validation

Within each cohort, the logarithm of the peak signal was regressed on the logarithm of the resting signal and the standardized residual taken as the conditioned response. Discrimination of subsequent fate was measured by the area under the ROC curve. Internal–external validation withheld one cohort at a time, with the baseline regression, the standardization and the classifier coefficients estimated only in the remaining cohorts and applied to the held-out cohort without re-estimation. The null distribution was built by permuting the conditioned response within cohort and within quintile of resting signal, 5,000 times, which preserves any information carried by the baseline itself. Timing of death was analysed by Cox proportional hazards with Breslow handling of ties and permutation inference.

### Bliss-independence comparison

Deviation from Bliss independence was calculated descriptively [45]. Survival was measured at defined times for each agent alone and in combination, and observed combination survival was compared with the value expected if the two interventions acted independently. The score is log₂(experimental survival/theoretical survival), so negative values indicate survival below the Bliss expectation and positive values indicate survival above it. Each arm was scored against controls acquired in the same experiment. Because the combination experiments contained single determinations per arm, these scores are descriptive and are not used as inferential evidence of reproducible synergy.

## Declarations

### Ethics approval

AML provide the archival human tumour material that was handled in accordance with the Declaration of Helsinki.

## Author contributions

F.J.I. conceived and designed the study, performed the imaging experiments and wrote the manuscript. R.G. and J.D.-C. performed the quantitative and kinetic analysis. A.M.-L. and J.L.G. contributed to the cell biology experiments. A.C. provided and characterized the human tumour material. P.M.I. and J.C. contributed to the design of the therapeutic experiments and to the interpretation of the results. All authors read and approved the manuscript.

## Data availability

The single-cell fluorescence trajectories, fate annotations and derived parameters supporting this study are available from the corresponding author on reasonable request and will be deposited in a public repository. Repository details and a resolvable identifier will be added to the preprint record when available.

## Acknowledgements

We thank the microscopy facilities of the Centro de Investigación Príncipe Felipe and the Instituto de Biomedicina de Valencia.

---

Figure S1. **PKM2 expression in tumour and healthy tissue.** PKM2 levels in tumour tissue (orange) compared with the corresponding healthy tissue (grey), from publicly available datasets through the GEPIA platform. BRCA, breast invasive carcinoma; CESC, cervical squamous cell carcinoma and endocervical adenocarcinoma; CHOL, cholangiocarcinoma; COAD, colon carcinoma; DLBC, diffuse large B-cell lymphoma; KIRP, kidney renal papillary cell carcinoma; LIHC, liver hepatocellular carcinoma; LUSC, lung squamous cell carcinoma; OV, ovarian serous cystadenocarcinoma; READ, rectum adenocarcinoma; SKCM, skin cutaneous melanoma; STAD, stomach adenocarcinoma; UCS, uterine carcinosarcoma; PAAD, pancreatic adenocarcinoma.

**Supplementary movies.** TMRM Cont shows the evolution of the TMRM signal through the experiment. Cells were incubated for 30 min with 500 nM TMRM and 10 ng/mL MitoTracker Green and imaged by fluorescence microscopy at 10 s intervals. MG Cont shows the same cells in the MitoTracker Green channel. As the TMRM signal declines the MG signal rises, which is the inverse relation described in Figure 5e,f.

